# Accurate Determination of Human CPR Conformational Equilibrium by smFRET using Dual Orthogonal Non-Canonical Amino Acid Labeling

**DOI:** 10.1101/407684

**Authors:** Robert B. Quast, Fataneh Fatemi, Michel Kranendonk, Emmanuel Margeat, Gilles Truan

## Abstract

Conjugation of fluorescent dyes to proteins - a prerequisite for the study of conformational dynamics by single molecule Förster resonance energy transfer (smFRET) - can lead to substantial changes of the dye’s photophysical properties, ultimately biasing the quantitative determination of inter-dye distances. In particular the popular cyanine dyes and their derivatives, which are by far the most used dyes in smFRET experiments, exhibit such behavior. To overcome this, a general strategy to site-specifically equip proteins with FRET pairs by chemo-selective reactions using two distinct non-canonical amino acids simultaneously incorporated through genetic code expansion in *Escherichia coli* was developed. Applied to human NADPH-cytochrome P450 reductase (CPR), the importance of homogenously labeled samples for accurate determination of FRET efficiencies was demonstrated. Furthermore, the effect of NADP+ on the ionic strength dependent modulation of the conformational equilibrium of CPR was unveiled. Given its generality and accuracy, the presented methodology establishes a new benchmark to decipher complex molecular dynamics on single molecules.

---

Numerous biophysical methods can be used to analyze proteins and derive detailed structural information. Albeit their immense power for supporting structural biology, most of these approaches do not resolve the structural dynamics occurring in individual molecules under physiological conditions, a prerequisite for linking conformational information to biochemical function^1^. In particular, conformational transitions may be averaged out using ensemble-based methods, thereby impairing the elucidation of potential short-lived intermediate conformations. For example, large structural reorientations occurring below the ms timescale have been observed for the human NADPH-cytochrome P450 reductase (CPR) using a variety of different biophysical methods.^2^-^6^ CPR is a paradigmatic diflavin reductase, localized at the endoplasmic reticulum, where it transfers electrons from NADPH to a variety of acceptors including P450 enzymes and plays a major role in essential physiological processes such as drug, steroid and fatty acid metabolism.^7^ CPR alternates between a locked, compact state, competent for electron transfer between its two flavin cofactors, bound to distinct FMN and FAD/NADPH domains, and an unlocked, open state, allowing electron transfer to acceptors (Figure 1). In the unlocked state, CPR has been described to experience a large relative conformational freedom of the two flavin binding domains around its flexible hinge segment.^5^ The first evidences of these structural rearrangements were provided by crystal structures of a mutant lacking 4 residues in the hinge region^8^ and a human:yeast chimera.^9^

**Figure 1.**
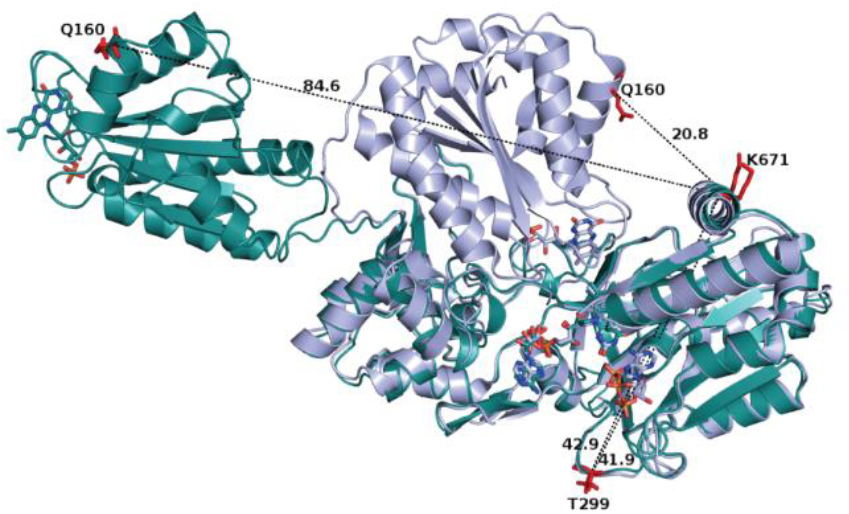
Overlay of compact (grey, PDB code 5FA6) and extended (green, human extended conformation model based on 3FJO) conformations of soluble human CPR with highlighted distances (Å) of residues selected for installation of fluorescent moieties (red). The FMN, FAD and NADP+ molecules are shown as sticks

Single molecule Förster resonance energy transfer (smFRET) allows for monitoring structural changes in proteins by reporting on distances in the range of 2-10 nm at nanosecond up to minute timescales.^1^ Nevertheless, smFRET measurements require the site-specific attachment of two organic dyes to the protein-of-interest. The photophysical properties of dyes are sensitive to their physicochemical environment, leading to altered quantum yields (QY) and fluorescence lifetimes,^10^ which in turn impact the determination of FRET efficiencies (for details see Supporting Information). However, these parameters can be experimentally determined, allowing for accurate calculation of transfer efficiencies and inter-dye distances provided that only one type of doubly labeled species is present in the sample.^11^ In contrast, such a precise correction remains impossible when proteins have been randomly labeled using only one chemical reaction, as each dye will have two different chemical environments affecting its photophysical properties.

Site-specific incorporation of ncAAs into proteins is possible using stop codon suppression mediated by orthogonal translation systems (OTS)^12^-^15^ to provide unique reactive centers for the selective attachment of dyes. Recently, Sadoine et al. labeled calmodulin by a combination of cysteine-maleimide coupling and bioorthogonal reaction of a ncAA to demonstrate the effect of the labeling position on FRET efficiencies.^11^ Here, we present a strategy for the accurate assessment of conformational dynamics in proteins by smFRET, exemplified on the soluble form of human CPR. As human CPR contains multiple surface exposed cysteines, one of them being critical for catalysis,^16^ we decided to employ a more general labeling strategy comprising two uniquely reactive non-canonical amino acids (ncAAs) followed by subsequent selective fluorescent modification using two bioorthogonal chemistries.

In contrast to previous studies,^14^ cotranslational incorporation of p-propargyloxy-L-phenylalanine (PrF) and cyclopropene-L-lysine (CpK), mediated by a mutant *M. janaschii* tyrosyl-tRNA synthetase (PrFRS)/ tRNA_CUA_^pair17^ and the wild type *M. barkeri* pyrrolysyl-tRNA synthetase (CpKRS)/ *M. mazei* tRNA_UUA_, is achieved through parallel suppression of premature *Amber* (TAG/UAG) and *Ochre* (TAA/UAA) stop codons (Figure 2a) using a single newly developed suppression vector. We further measured the influence of the dye position on QYs and demonstrate that our labeling strategy enables accurate determination of FRET efficiencies. Finally, we modulate the conformational dynamics of CPR by salt and confirm the role of the oxidized substrate NADP+ in stabilizing the locked state of CPR.

**Figure 2.**
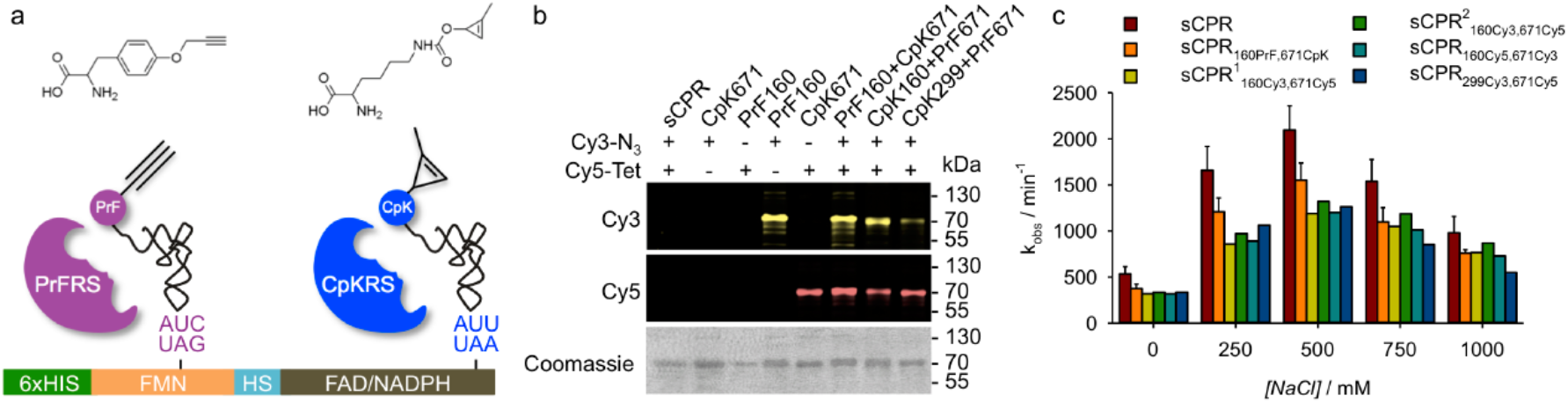
Cotranslational incorporation of ncAAs and chemoselective fluorescence modification. a) Parallel incorporation of PrF (left) by Amber (purple) and CpK (right) by Ochre (blue) suppression mediated by PrFRS/tRNACuA and CpKRS/tRNAUUA, respectively. The schematic representation of the sCPR gene includes the N-terminal 6 x Histidine tag (green), the FMN domain (yellow), the hinge segment (HS, cyan) and the FAD/NADPH domain (brown). b) Bioorthogonal modification of incorporated PrF with Sulfo-Cy3-Azide (Cy3-N3) by CuAAC and CpK with Tetrazine-Sulfo-Cy5 (Cy5-Tet) by SPIEDAC. c) Cytochrome c reduction efficiency of labeled sCPR variants given by the initial velocity kobs. Error bars represent the standard deviation of triplicates. For sCPR_160Cy3,671Cy5_ data is shown for individual samples later used in smFRET measurements.

Based on structures of CPR in a locked (PDB ID 5FA6) and a potential maximally extended state (derived from PDB ID 3FJO, see methods section), surface exposed residues Q160 in the FMN and K671 in the FAD domain were selected for incorporation of ncAAs (Figure 1). Inter residue distances were estimated to be ∼30 Å and ∼85 Å for the two conformations, leading to a large FRET efficiency change given the reported Forster radius of 50-55 Å for the Cy3:Cy5 pair.^18^ Additionally, T299 served as a constant FRET control in combination with K671, as no change in the inter-residue distance was predicted during the FMN domain movement.

Initial attempts to utilize the previously developed pUltraII system^12^ for co-expression of the two OTSs failed to produce soluble sCPR (data not shown). Thus, a novel vector was designed (pRQ101), harboring one cassette for each of the OTSs constructed on the basis of the previously described enhanced pEVOL system.^19^ This novel suppression vector allows for an independent induction of the synthetases, thereby enabling a gentle expression of sCPR, obtained through transcriptional leakage of the *lac* promoter in the absence of IPTG. In this way, a set of functional sCPR variants harboring one or both ncAAs at varying positions was produced at yields comparable to previous studies^14^ (1-4 mg/L of purified protein for double incorporation depending on the position of premature stop codons (Supplementary Figure S1), and separated from the truncated termination products by two consecutive affinity purifications (Supplementary Figure S2). Although fluorescence modification in a one-pot reaction failed due to unwanted side reactions (Supplementary Figure S3), sequential labeling of CpK by inverse electron-demand Diels-Alder cycloaddition (SPIEDAC) with 6-Methyl-Tetrazine-Sulfo-Cy5 (Cy5-Tet) followed by Copper(II)-dependent azide-alkyne cycloaddition (CuAAC) of PrF with Sulfo-Cy3-Azide (Cy3-N3) provided perfect specificity (Figure 2b and Supplementary Figure S3) and satisfying efficiency under optimized conditions (Supplementary Figure S4, S5, S6). The degree-of-labeling (DOL) for the doubly labeled variants estimated from the corrected absorbance of dyes and bound flavin cofactors was generally >85 % for SPIEDAC and >55 % for the subsequent CuAAC. Accordingly, determination of the stoichiometry of donor and acceptor dyes on measured single molecules further confirmed the presence of about 1/3 of doubly labeled molecules (data not shown). In addition, the labeled samples of sCPR showed little differences in electron transfer activity (Figure 2c), that were consistent within all doubly labeled constructs and similar to batch-to-batch differences routinely observed between biological replicates.^920^ Even more importantly, the functional behavior of labeled sCPR variants in response to different NaCl concentrations with a maximal activity at ∼500 mM was retained (Figure 2c and Supplementary Figure S7).

The photophysical behavior of the popular cyanine dyes Cy3 and Cy5 is known to be influenced by their local environment.^10^ Accordingly, we generated two variants of sCPR in which the positions of donor and acceptor dyes were reversed (Figure 3, sCPR_160Cy3,671Cy5_ and sCPR_160Cy5,671Cy3_) and evaluated the establishedFRET system by collecting data of freely diffusing single molecules on a homemade confocal setup using alternating pulsed laser excitation (ALEX/PIE).^21^

**Figure 3.**
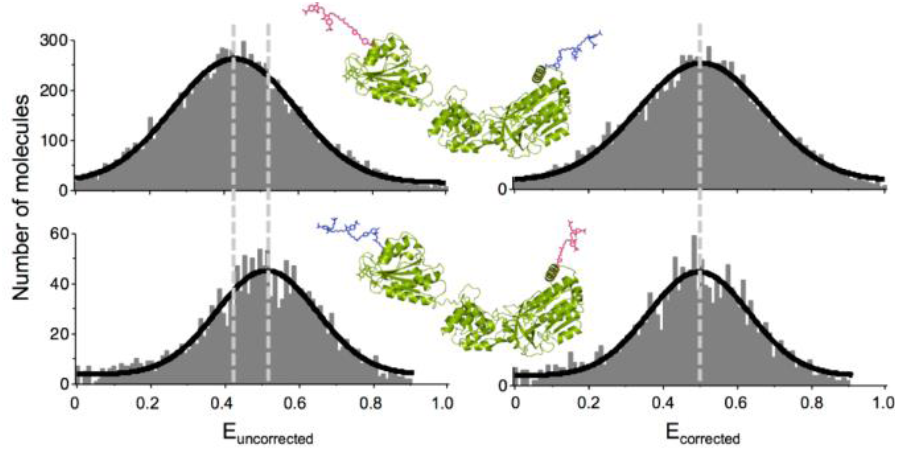
Influence of labeling position on FRET efficiency. uncorrected (left) and QY corrected (right) histograms of sCPR labeled with Cy3 (magenta) on FMN and Cy5 (blue) on FAD domain (sCPR_160Cy3,671Cy5_ top) and vice versa (sCPR_160Cy5,671 Cy3_ bottom). Dashed lines indicate peak positions.

Distance determination in smFRET experiments depends on the correct determination of the fluorophores quantum yields (Φ_F_). Indeed, the D A distance (*R*) is calculated in a FRET experiment using the following equation, where two terms depend on Φ_F_:

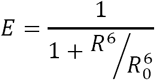

Firstly, the Forster radius *R*_*o*_ depends on the donor QY ϕ_F,D_ as follows:

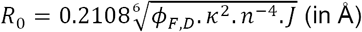

where *κ*^*2*^ is the orientation factor, *n* the medium index of refraction and *J* the spectral overlap integral (in cm^-1^M^-1^nm^4^).

Secondly, the transfer efficiency *E* is measured in a ratiometric smFRET experiment using:

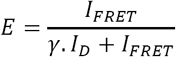

where *I*_*D*_ and *I*_*FRET*_ represent the fluorescence intensities detected for each molecule, i.e. donor and acceptor emissions, respectively (corrected for background and other crosstalk parameters, see Materials and Methods section for details) and where γ represents a correction factor that takes into account the fluorescence QYs of both dyes, and the donor *(G*_*G*_|_D_) and acceptor (*G*_*R* |*A*_)detection efficiencies (which depend on the instrumental characteristics and the emission spectral properties of the fluorophores).

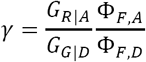

Thus, any change in the fluorescence QYs (Φf) of the dyes needs to be accounted for, to accurately calculate *γ, E, R_O_* and thus R. In this context, it is important to use a homogeneous sample where Φ_F,D_ and Φ_F,A_ can be measured for a specific position at which the dye is attached to the protein, a condition that is not fulfilled when using D-A statistical labeling on two cysteines for example. This requirement is particularly important when using cyanine dyes (such as Cy3, Cy5, and their derivatives), that are widely used in single molecule FRET experiments, and for which local environmental properties lead to strong changes in QYs and fluorescence lifetimes, notably as a consequence of modulation of the rate at which the molecules isomerize from the photo-active trans into the dark cis state^10,22,23^.

Often, the apparent FRET efficiency (or proximity ratio) *E_PR_* is reported in the literature when only relative distance changes are considered and is calculated without taking into account the γ correction factor:

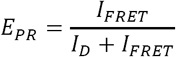

Here, comparison of the apparent FRET efficiency *E*_*PR (uncorrected)*_ histogram measured for sCPR_160Cy3,671 Cy5_, and sCPR_160Cy5,671 Cy3_, in the unlocked (open) state, strikingly demonstrated the effect of dye quantum yield changes due to their local environment on the calculation of this parameter (Figure 3). However, thanks to our orthogonal labeling strategy, we generated four singly labeled constructs and calculated the individual, position specific, donor and acceptor quantum yields (Supplementary Table S1). Accounting for these differences, the corrected FRET efficiency distributions turned out to be nearly identical for sCPR_160Cy3,671Cy5_, and sCPR_160Cy5,671 Cy3_,, underlining the value of the presented approach for acquisition of accurate smFRET data (Figure 3, E_corrected_).

Previous work had shown that the efficiency of cytochrome *c* reduction by sCPR is directly related to the conformational equilibrium, which can be modulated by ionic strength.^35^ Accordingly, the FRET efficiencies of the sCPR_160Cy3,671 Cy5_ variant decreased with increasing concentrations of NaCl (Figure 4a). This corresponds to an increase of the average inter-dye distance and thus a shift of the equilibrium in favor of the unlocked state. We validated this interpretation using two constructs. First, the sCPR_160Cy5,671 Cy3_, for which the two dyes are inverted, displayed a similar salt-dependent shift of the FRET distribution as expected (data not shown). Second, the aforementioned sCPR_299Cy3,671 Cy5_ variant showed no inter-dye distance changes in response to ionic strength variations (Supplementary Table S2 and Figure S8).

**Figure 4.**
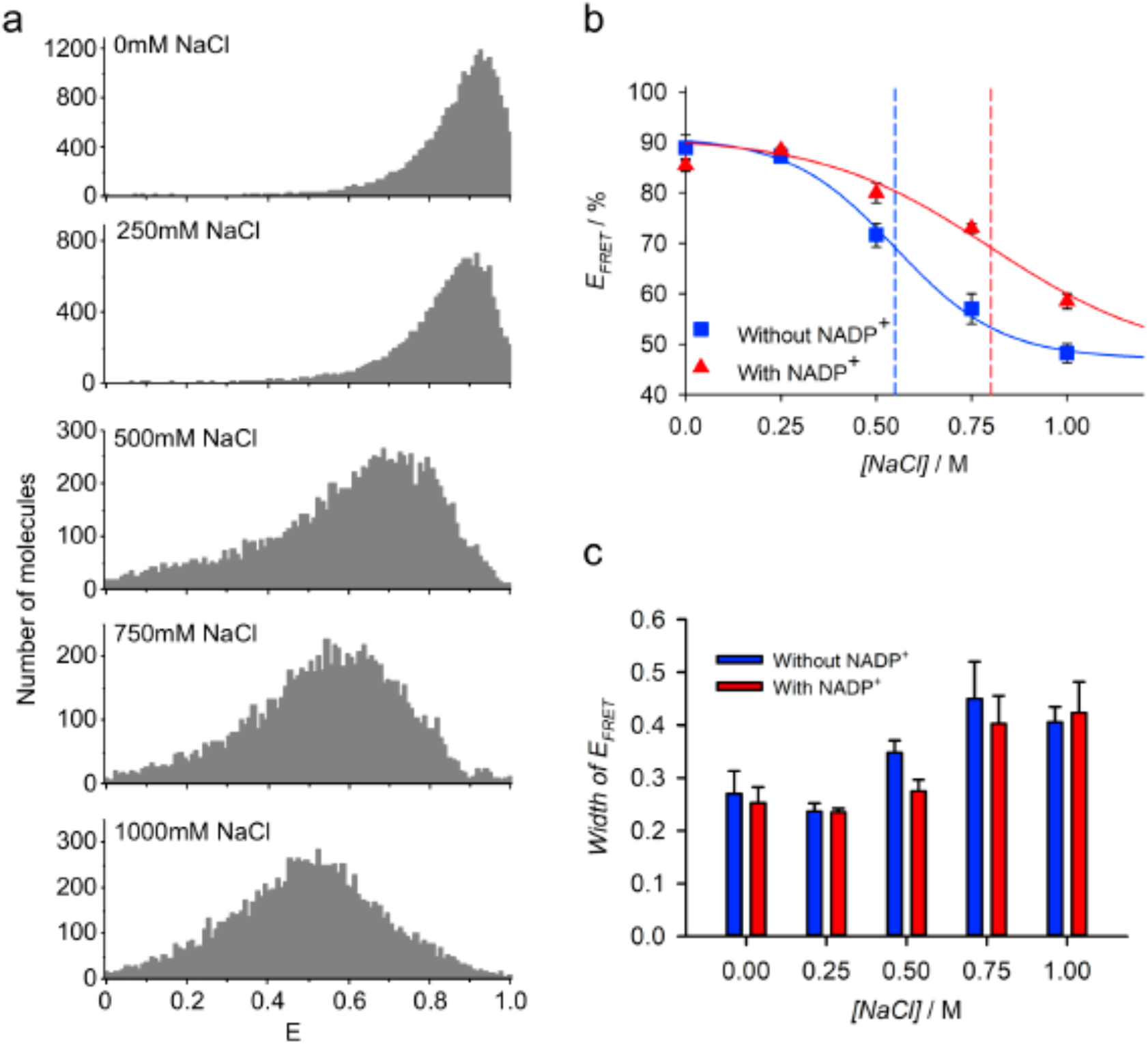
Ionic strength dependent modulation of conformational equilibrium between locked and unlocked states. a) FRET efficiency histograms of sCPR160Cy3,671Cy5 at different NaCl concentrations. b) Influence of NADP+ on equilibrium. Data points represent maxima of the FRET efficiency in the absence (blue squares) and presence (red triangles) of 500 μM NADP+. Functions were derived by fitting a two state model (see text) and the mid-point transition is indicated by a dashed line. c) Widths of FRET efficiency distributions fitted by a Gaussian function, at different NaCl concentrations.

We note that in all cases, the width of the FRET efficiency distributions were wider than what would be predicted for a single donor-acceptor distance, taking into account the distribution broadening due to photon shot noise. This effect is particularly visible at conditions above physiological ionic strength (≥500 mM), and could point to the presence of multiple conformations in the unlocked state, interconverting in the ms timescale, in accordance with previous SAXS, NMR^5^ and smFRET data.^24^

Several studies have provided evidence for a closing of sCPR induced by its oxidized substrate NADP+.^2,25,26^ For comparison we fitted the smFRET data obtained in the presence and absence of 500 pM NADP^+^ to a previously developed salt-dependent two state model of the conformational equilibrium.^5^ In the presence of NADP^+^ a shift of the mid-point transition to higher NaCl was observed (Figure 4b). However, the overall distribution of FRET efficiencies remained strikingly similar when compared to data obtained in the absence of NADP+ (Figure 4c). Notably, a maximal average distance of 57 Å of the two dyes was calculated in the unlocked state at 1 M NaCl, which was only slightly reduced to 53 Å when NADP+ was present. Altogether, NADP+ binding seems not to be sufficient to close the sCPR structure per se, but rather shifts the mid-point of the conformational equilibrium to stabilize the locked state and has no general effect on the distribution of potential conformational subpopulations. Physiologically, the ionic strength is equivalent to about 170 mM NaCl, and the reported NADP^+^ concentration is around 5 µM (100 times lower than the one we used).^27^ We can thus hypothesize that the shift in the equilibrium induced by the oxidized cofactor on oxidized CPR is probably minor under physiological conditions.

In this study, we presented a general strategy for the domain-selective equipment of human CPR with a FRET pair to enable an accurate single molecule analysis of conformational dynamics. The developed labeling strategy is of high value for attachment of dyes to any protein for which cysteine and other labeling strategies are inapplicable. It is easily adaptable for the study of diverse biological questions because i) the used OTSs, ncAAs and the FRET pair are easily accessible from academic laboratories and/or commercial sources, ii) they are interchangeable allowing for free choice of ncAAs, labeling reactions and FRET pair, iii) only a single suppression vector has to be co-transformed together with the expression vector for the protein-of-interest and iv) the parallel suppression of two stop codons eliminates the need for any additional components to facilitate efficient ncAA incorporation with high fidelity.

Although a variety of biophysical methods have been applied to study the domain movements in CPR,^2,45,25,26,28,29^ the newly established FRET system offers a valuable complementation as it enables studying single molecules at timescales inaccessible by other methods within complex biological environments. With the fast pace of technical and methodological advances we envision that this system will provide new insights into the structural dynamics of CPR as a prototypical diflavin reductase but also help to assess, with a very high precision, the potential roles of cofactor binding on structural transitions. As such, CPR can be considered as a model for multidomain proteins exhibiting large and fast conformational transitions.

## ASSOCIATED CONTENT

A detailed experimental description of template generation, protein expression, protein purification, protein labeling, cytochrome c reduction assay, quantum yield determination, smFRET measurements and data analysis, modeling of CPR open conformation, curve fitting, calculation of Förster radius and total distances, supplementary figures, supplementary tables and gene and protein sequences is available in supplementart material

## ACKNOWLEDGMENTS

This work was in part funded by a joint ANR/FCT program; France: ANR-13-ISV5-0001 (DODYCOEL), and Portuguese national funds, through the Funda?ao para a Ciencia e a Tecnologia (Project FCT-ANR/BEX-BCM/0002/2013). The CBS belongs to the France-BioImaging infrastructure supported by the French National Research Agency (ANR-10- INBS-04, " Investments for the future"), and is supported by the GIS " IBiSA: Infrastructures en Biologie Sante et Agronomie". Additonally, RBQ was supported by an " Access grant" from the France-BioImaging infrastructure. We thank the committee of the INSEERE project, in particular Regis Faure, Sebastien Nouaille and Virginie Ramillon-Delvolve for assistance in molecular biology and Hélène Cordier for access to the Biorad Imaging System. Further we thank Philippe Urban for critical reading of the manuscript.

## Supporting Information

## Materials and Methods

### Materials

All chemicals were purchased from Sigma (France) and Roth (France) if not stated otherwise. Enzymes used for molecular biology were purchased from Thermo Fischer (France).

### Template generation

Generation of the expression vector pET15b harboring the non-codon optimized gene of the soluble form of human CPR, which lacks the N-terminal 44 amino acid coding region, and contains a N-terminal 6xHis-tag followed by a thrombin cleavage site, was described previously ^1^. Mutations were introduced by PCR amplification of the complete vector using completely complementary primer pairs harboring the desired substitutions or 5’ phosphorylated primers generated using the NEBaseChanger software (New England Biolabs). All primer sequences can be found in S10 Table. Initially, the TAG *Amber* codon terminating the TAG ORF was substituted to the TGA ***Opal*** codon using primers 1/2. Subsequently, this template DNA was used to generate CPR mutants containing premature TAG and TAA codons.

The vector pRQ001 used for co-expression of MyPrFRS/tRNA_CUA_ and MbCpKRS/ MmtRNA_UUA_ was generated in several steps based on the previously described enhanced pEVOL system ^2^ Firstly, the MbPylRS gene in the pUltraII vector (gift from PG Schultz, Scripps Institute, La Jolla, USA), containing an Y349F mutation was mutated back to the wild type sequence (UniProtKB - Q6WRH6) using primers 11/12 as described above. Secondly, the wild type PylRS gene was amplified by PCR using primers 13/14 and 15/16 in order to introduce *BglII/SalI* and *NdeI/PstI* restriction sites upstream/downstream of the ORF, respectively. The pEvol pylRSWT (containing the wild type *M. mazei* PylRS [UniProtKB - Q8PWY1], gift from EA Lemke, EMBL Heidelberg, Germany) as well as the PCR products were cut accordingly to integrate the amplified wild type MbPylRS genes by replacing the *M. mazei* PylRS genes. Subsequently, the pyrrolysyl-tRNACUA gene of the newly generated pEVOL-MbPylRS_CUA_ was mutated to change the anticodon CUA to UUA with primers 17/18, enabling suppression of the UAG *Amber* codon. In the same step, a U25C substitution was introduced, which has previously been described to increase the suppression efficiency ^3^. This new vector designated pEVOL-MbPylRS_UUA_ was cut with *XhoI* and treated with the Klenow exonuclease fragment (New England Biolabs, France) to generate blunt ends. A pEVOL vector containing two copies of the PrF-specific mutant of *Mj* tyrosyl-tRNA synthetase ^4^ (Y32A, E107P, D158A, L159I, L162A, D286R, gift from Régis Fauré, LISBP, Toulouse, France) as well as the corresponding *Amber* suppressor tRNA_CUA_ was cut with *XhoI/BamHI* and likewise treated to obtain blunt ends. Finally, the fragment containing the MjPrFRS/tRNA_CUA_ was ligated into the pEVOL-MbPylRS_UUA_ to give pRQ101. Gene sequences of sCPR, synthetases and tRNAs can be found in S11 Gene Sequences.

### Protein expression and purification

*E. coli* BL21 (DE3) cells were co-transformed with pRQ101 and pET15b-CPR and selected on LB-Agar containing 30 μg/ml Chloramphenicol and 50 μg/ml Ampicillin. A 10 ml pre-culture was inoculated from a single colony and grown over night at 37°C in LB medium supplemented with 30 μg/ml Chloramphenicol and 50 μg/ml Ampicillin. The grown pre-culture was used to inoculate 200 ml TB medium at a 1:100 ratio in a 1 L Erlenmeyer flask supplemented with 30 μg/ml Chloramphenicol and 50 μg/ml Ampicillin, 100 μM Riboflavin, 2 mM MgSO_4_ and 1 mM of PrF (Iris Biotech, Germany), CpK (Sirius Fine Chemicals, Germany) or both. The culture was grown at 28°C without addition of IPTG in order to take advantage of the leakage of the ***lac*** operon promoting a slow expression of CPR and thereby providing soluble, properly folded and functional sCPR. After 4-6 h culturing, L-arabinose was added to a final concentration of 0.02%. Cells were harvested by centrifugation at 6000 x g and 4°C for 10 min following a total of 48 h post-inoculation. Cell pellets were washed with PBS and subsequently resuspended in lysis buffer (20 mM Na/K PO_4_, pH 7.4, 250 mM NaCl, 250 mM KCl, 1 x protease inhibitor cocktail (0.3 μM aprotinin, 1 μM leupeptin, 1.5 μM pepstatin A, 100 μM benzamidine, 100 μM metabisulfite), 1mM Tris-(2-carboxyethyl)-phosphin, 2 μM FMN, 2 μM FAD, Lysozyme and DNaseI). The resuspended cells were incubated for 30 min at RT and disrupted by 4 consecutive freeze-thaw cycles using liquid N2 and a water bath at 42°C with rigorous vortexing in between. Subsequently, the soluble fraction was obtained by centrifugation at 10.000 x g and 4°C for 1h. For an initial purification sCPR containing a N-terminal 6xHis-tag was bound to TALON Superflow resin (GE Healthcare, France). After washing with a 20 mM Na/K PO_4_ (pH 7.4) buffer containing 250 mM NaCl, 250 mM KCl, 2 pM FMN and 2 pM FAD, sCPR was eluted with 20 mM Na/K PO4, (pH 7.4) containing 250 mM NaCl, 250 mM KCl, 2 pM FMN, 2 pM FAD and 250 mM Imidazole. Protein was oxidized with potassium ferricyanide prior to a concentration step using Amicon Ultra 0.5 mL centrifugal filters (cut-off of 50 kDa (Merck, France)) with a centrifugation at 14000 x g and 4°C for 5-10 min. Finally, five consecutive rounds of concentration and dilution with 20 mM Tris-HCl, pH 7.4 (approximately 10 to 20-fold per dilution) were used to remove residual traces of imidazole and exchange the buffer. A second purification step was performed using 2’5’-ADP-Sepharose 4B (GE Healthcare, France). The 6xHis-tag purified sCPR was diluted approximately 1:1 with 20 mM Tris-HCl (pH 7.4) containing 150 mM NaCl and bound to the resin, followed by washing with the same buffer. The protein was eluted with a 20 mM Tris-HCl (pH 7.4) buffer solution containing 500 mM NaCl, 2 μM FMN, 2 μM FAD and 1 mM NADP+. Subsequent concentration and buffer exchange were performed using Amicon centrifugal filters as described above. The sCPR concentration was determined by absorption spectroscopy based on the known absorbance maximum of the oxidized flavin cofactors (ε_454nm_ = 22,000 cm^-1^M^-1^). Protein integrity was verified by SDS-PAGE, Western Blot (1^st^ antibody: 6x-His Tag Monoclonal Antibody (HIS.H8), MA1-21315, Thermo Fischer, France; 2^nd^ antibody: Goat anti-Mouse IgG (H+L) Secondary Antibody, HRP, #31430, Thermo Fischer, France; detection: SIGMAFAST BCIP/NBT, Sigma, France). Protein functionality was verified by determination of the cytochrome ***c*** reduction efficiency k_obs_ (see below).

### Protein labeling

Typical labeling reactions were performed in a total volume of 50-100 μl at a final concentration of 20 μM sCPR. The strain-promoted inverse electron-demand Diels-Alder cycloaddition (SPIEDAC) was carried out at 30°C for 16 h in 20 mM Na/K PO_4_ buffer (pH7.4) containing 500 mM NaCl and 200 μM 6-Methyl-Tetrazine-Sulfo-Cy5 (Jena Bioscience, Germany). Following incubation, the reaction was diluted to 500 μl with a 20 mM Na/K PO_4_ buffer (pH 7.4) containing 2 μM FMN, 2 μ M FAD and the sample was subsequently concentrated using Amicon centrifugal filters (as described above). This wash and concentration procedure was repeated three times to remove the majority of free Cy5 dye, followed by a final gel filtration step using Zeba Spin Desalting Columns, 40K MWCO, 0.5 mL (Thermo Fischer, France) equilibrated with the same buffer but lacking FMN and FAD.

For the copper(I)-dependent alkyne-azide cycloaddition (CuAAC), first a 20 x stock solution containing 10 mM CuSO_4_ and 50 mM THPTA (Tris(3-hydroxypropyltriazolylmethyl)amine) was prepared in 20 mM Na/K PO_4_ buffer (pH7.4). A 50 μl reaction mixture was set up by mixing 2.5 pl of the CuSO4:THPTA stock with 20 mM Na/K PO_4_ buffer (pH7.4). Subsequently, sCPR was added (20 μM final concentration) followed by Sulfo-Cy3-Azide (Jena Bioscience, Germany) at a final concentration of 200 μM. Finally, the reaction was started by addition of 2.5 μl of a 100 mM stock of freshly prepared sodium L-ascorbate and quenched after one minute by addition of 50 μl cold 20 mM Na/K PO_4_ buffer (pH7.4) containing 2 mM EDTA. Dye removal was achieved as described above for the SPIEDAC labeling.

Labeling specificity and efficiency were estimate by use of absorption spectroscopy and in-gel (SDS-PAGE) fluorescence using a ChemiDoc MP Imaging System (Biorad). The labelling efficiency was estimated, based on the absorbance maxima of the dyes provided by the manufacturer (ε _647nm_ 6-Methyl-Tetrazine-Sulfo-Cy5 = 251,000 cm^-1^.M^-1^; ε_553nm_ Sulfo-Cy3-Azide = 151,000 cm^-1^.M^-1^). Concentrations of labeled sCPR were calculated based on values of oxidized flavin cofactor absorbance at 454 nm after spectral deconvolution using a homemade software (with kind permission of Denis Pompon).

### Cytochrome *c* reduction assay

The activity of sCPR and labeled variants thereof was assessed at salt concentrations corresponding to the ones used for smFRET measurements by determining the initial velocity of cytochrome ***c*** reduction referred to as *k***_*obs*_** A typical reaction was performed in a plastic cuvette in a total volume of 1 ml at 22-26°C. First, 943 μl of 20 mM Tris-HCl at pH7.4 containing either 0, 250, 500, 750 or 1000 mM NaCl were mixed with 50 μl 2 mM cytochrome ***c*** (#C3131 Sigma, France), followed by addition of 2 εl 100 mM freshly prepared NADPH. After thorough but careful mixing the acquisition was started to record the baseline and upon addition of 5 εl sCPR at 2 εM the reduction of cytochrome ***c*** was recorded at 550 nm with 0.1 sec averaging for a total of 2 min using the Varian Cary 100 Scan UV-Vis spectrophotometer (Agilent, France). The initial velocity kobs was calculated using the slope of the linear part of the curve and ∇ ε _550nm_ = 21,000 cm^-1^.M^-1^ for reduced cytochrome *c*.

### Quantum yield determination

Relative quantum yields (QY) for singly labeled constructs were determined by the so-called comparative method^5^ using the Varian Cary spectrophotometer and the SAFAS Xenius XC spectrofluorimeter (SAFAS Monaco, Monaco). After labeling and purification, singly labeled sCPR was diluted to obtain a maximal OD of 0.1 at wavelength higher than the excitation wavelength in buffer containing non-labeled sCPR to give a total final concentration of labeled + non-labeled sCPR of 1 μ M, preventing dilution-dependent flavin cofactor loss. Absorbance and fluorescence spectra were recorded for 5 dilutions at 22-26°C and QY were calculated using cresyl violet (QY 0.54 in methanol) and Cy5 (QY 0.27 in PBS) as standards for Sulfo-Cy3-Azide (excitation 515 nm, emission 525 - 800 nm) and 6-Methyl-Tetrazine-Sulfo- Cy5 (excitation 600 nm, emission 610 - 800 nm), respectively.

### smFRET measurements and data analysis

smFRET measurements were performed on a home-built confocal PIE-MFD microscope as described previously^6^. Samples at a concentration of 30 to 100 pM labeled sCPR in 20 mM Tris-HCl (pH7.4) with 0, 250, 500, 750 or 1000 mM NaCl and supplemented with 1 μM non-labeled sCPR to prevent cofactor loss were placed in transparent non-binding 384-well plates (Corning, USA) and acquisitions were performed for at least 2 h at room temperature (22°C). The power of green and red laser beams was set to 25 μW each before entering the microscope. Collected data were analyzed with the " Software Package for Multiparameter Fluorescence Spectroscopy, Full Correlation and Multiparameter Fluorescence Imaging" developed in CAM Seidel’s laboratory (www.mpc.uni-duesseldorf.de)^7^. A single molecule event was defined as a burst containing at least 30 photons. For each photon burst, we calculated the intensity-derived donor-acceptor FRET efficiencies (E), and the apparent FRET efficiencies (E_PR_)^8^ using the recommendations of a recent worldwide benchmark study on precision and accuracy of single-molecule FRET measurements^9^. These calculations require the correction of the background from donor and acceptor channels, the factor for spectral crosstalk (α), arising from donor fluorescence leakage into the acceptor channel, the factor for direct excitation (δ) of the acceptor with the donor laser and the detection correction factor (γ). In our experimental configuration, we found α=0.07 and 5=0.07. E_PR_ was calculated as follows:

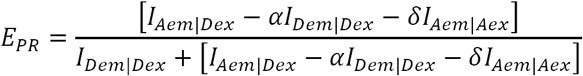

where *l*_*xΦ Y*_ are the background-corrected acceptor emission upon donor excitation *(I*_*Aem*_ *\Dex*), donor-emission upon donor excitation *(l***_*Dem*_** *\Dex*), and acceptor-emission upon acceptor excitation *(I*_*Aem*_ Φ _*Aex*_). E_PR_ does not take into consideration the detection correction factor (γ). The calculation of the FRET efficiency (E) thus requires the determination of the ratio of the donor (G_G Φ D_ p) and acceptor (G_R Φ A_) detection efficiencies (arXiv:1710.03807 [q-bio.QM]), and fluorescence quantum yields ***(Φ* _*F,D*_** and ***Φ* _*F,A*_ ).**

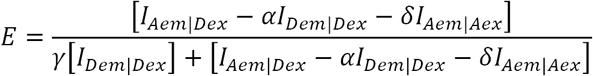

where,

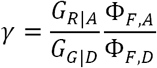

### Model of an open conformation of the human soluble CPR

The structure of the yeast:human chimera (PDB ID 3FJO)^1^ was used to create a model of the human soluble form of CPR in a potential extended conformation. An alignment between the yeast:human chimera and the human soluble form of CPR (sequence from PDB ID 5FA6^10^ was generated using CLUSTALW(http://www.genome.jp/tools-bin/clustalw). The YASARA software suite (YASARA Bioscience GmbH, Germany) was used to build the model using the AMBER03 force field.

### Curve fitting of FRET efficiencies vs. salt concentrations

The curve fitting was performed as described in^11^. Briefly, the FRET efficiency (E) was used to calculate the equilibrium constant ***K***_*e*_ between the locked and unlocked states of human soluble CPR with the following equation:

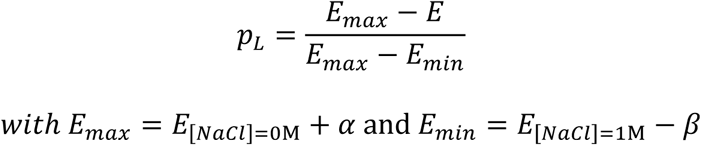

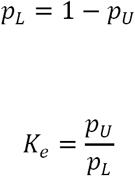

where ***p***_*U*_ represents the proportion of the unlocked state, ***pL*** the proportion of the locked state, ***E***_*max*_ the maximum FRET efficiency, ***E***_*min*_ the FRET efficiency at infinite NaCl concentration, α the difference between the maximum FRET efficiency and the one obtained at 0 M NaCl and β the difference between the FRET efficiency at 1 M NaCl and the one at infinite NaCl concentration. Obtained ***K*_*e*_** values where then fitted to the following equation with the Solver function of Excel (Microsoft, USA) using m, ∇G^w^, α and β as parameters.

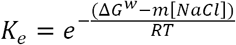

### Calculation of Förster radius R_0_ and total distances R

The apparent Förster radius ***R*_*0*_** for the Cy3-Cy5 FRET pair conjugated to sCPRi_160Cy3,671Cy5_ was calculated using the following equation

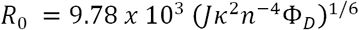

where ***J*** is the spectral overlap integral of the donor emission and acceptor excitation spectra, *κ*^*2*^is the orientation factor (2/3 in accordance with freely rotating dye molecules attached to the sCPR, which was supported by anisotropy measurements), ***n*** is the refractive index of the medium (1.4) and ***ϕ*_*D*_** the donor quantum yield (56.4 Å).

The calculated ***R*_*0*_** was then used to obtain total distances ***R*** according to

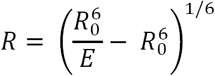

for the different experimentally determined peak values of the FRET efficiency ***E.***

## Supplementary Figures

**Figure S1.**
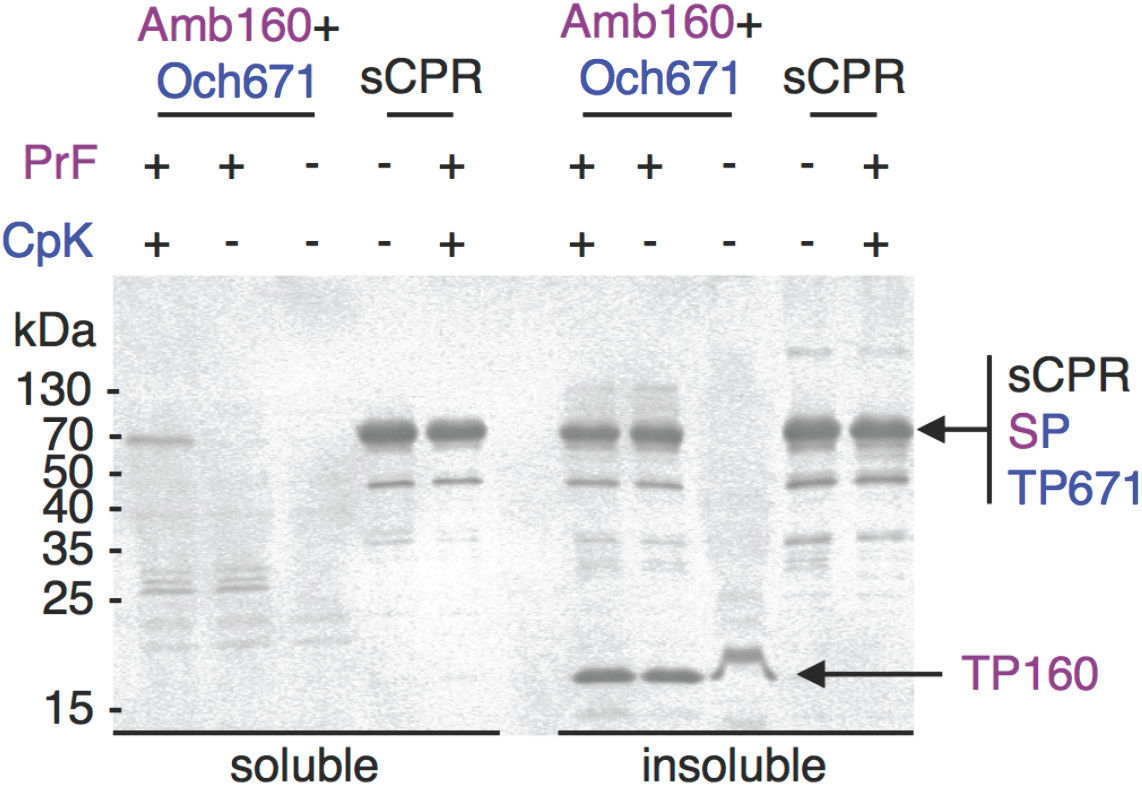
Site-directed incorporation of two distinct ncAAs in human sCPR. Immunoblot of soluble and insoluble fractions after lysis of cells expressing wild-type or sCPR_160Prf,671CpK_ gene harboring the two internal stop codons (Amb160+Och671) in the presence or absence of the non-canonical amino acids. Full-length sCPR, the suppression product (SP=sCPR_160Prf,671CpK_) and the two termination products at Amb160 (TP160) and Och671 (TP671) are marked by arrows. The incorporation of PrF and CpK is color coded in magenta and blue, respectively.

**Figure S2:**
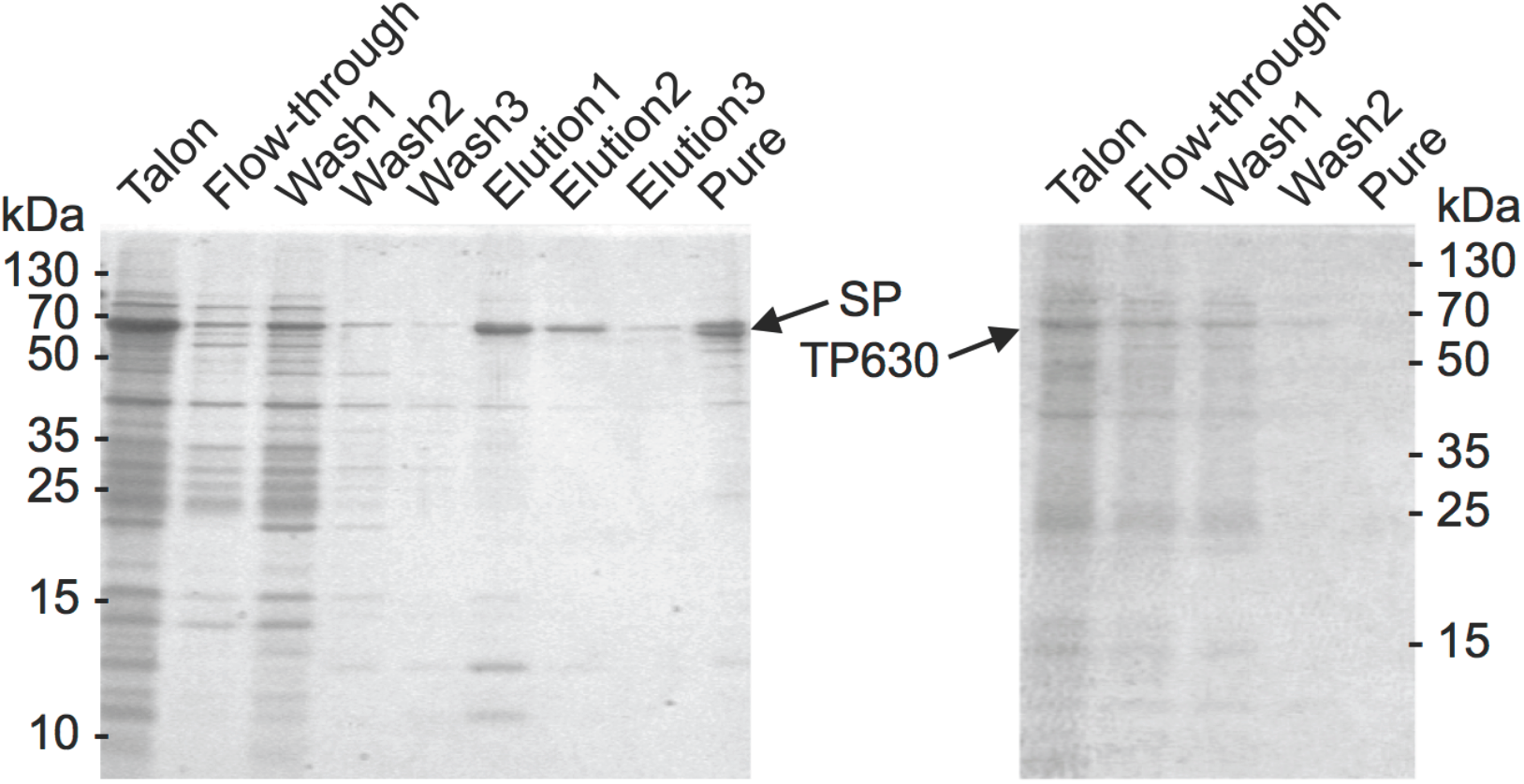
Selective purification of full-length sCPR suppression product (SP) by affinity chromatography using 2’5’-ADP sepharose. The full-length suppression product binds to the affinity matrix (left) while the truncated sCPR (TP630) lacking the 10 C-terminal amino acids does not bind and therefore is not present in the final concentrated eluate (right).

**Figure S3:**
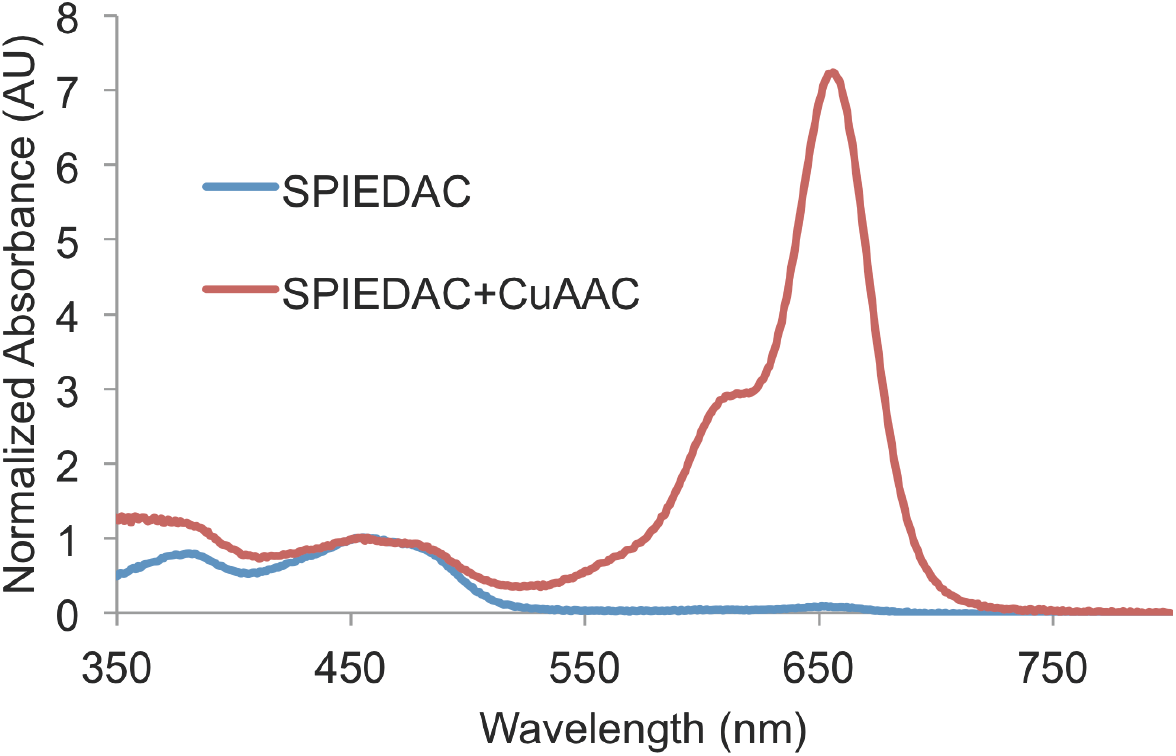
Incompatibility of SPIEDAC and CuAAC. The wild-type sCPR gene, not containing any premature stop codons was expressed in the presence of the orthogonal systems (PrFRS/tRNA_CUA_ and PylRS/tRNA_UUA_) and the ncAAs PrF and CpK, purified and subsequently treated with Tet-Cy5 only (SPIEDAC) or with Tet-Cy5, N_3_ -Cy3 and corresponding click reagents (CuSO_4_, Na+- Ascorbate, THPTA, SPIEDAC+CuAAC). The absorption spectra normalized to the flavin cofactor maximum at 454nm show that sCPR is modified by Tet-Cy5 only when being treated with the CuAAC reagents (red curve), which demonstrates an unspecific side reaction occurring under these conditions. In contrast, no labeling is observed when the wild type protein is treated only with Tet-Cy5 (blue curve), underlining the good specificity of the SPIEDAC reaction in the absence of the CuAAC reagents and further providing evidence that CpK is only incorporated in response to stop codon suppression in the mutant sCPR gene variants harboring premature stop codons.

**Figure S4:**
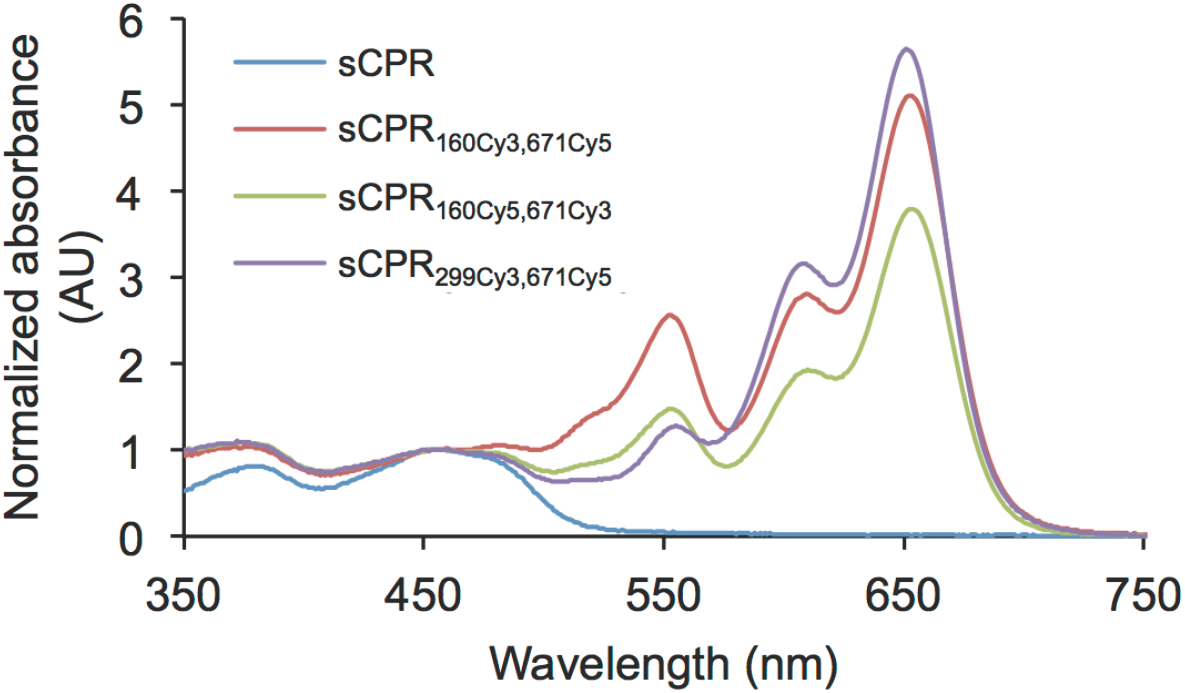
Absorption spectra of labeled sCPR variants. The absorption spectra were normalized to the flavin cofactor maximum at 454nm.

**Figure S5:**
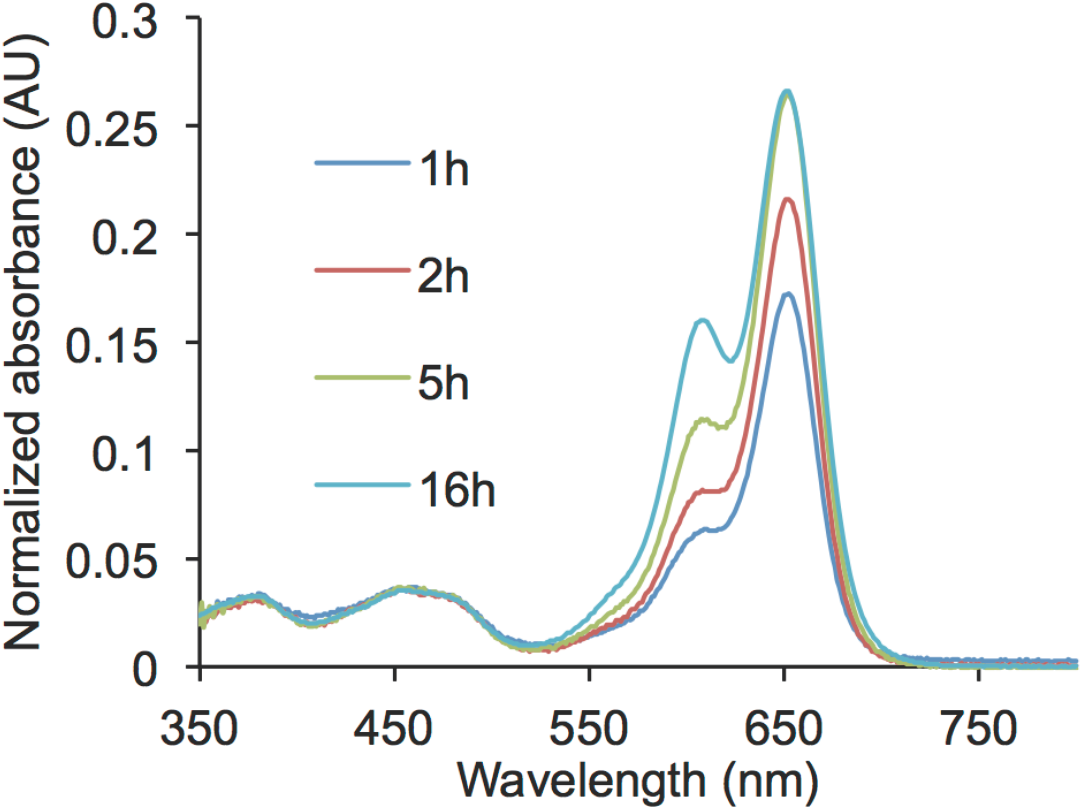
Time-course of the inverse electron-demand Diels Alder cycloaddition (SPIEDAC) reaction between sCPR_671CpK_ and 6-Methyl-Tetrazine- Sulfo-Cy5 at 30°C. Spectra were normalized to the flavin cofactor absorbance maximum at 454 nm and show that the degree-of-labeling reaches a plateau at 5h of reaction time at 30°C.

**Figure S6:**
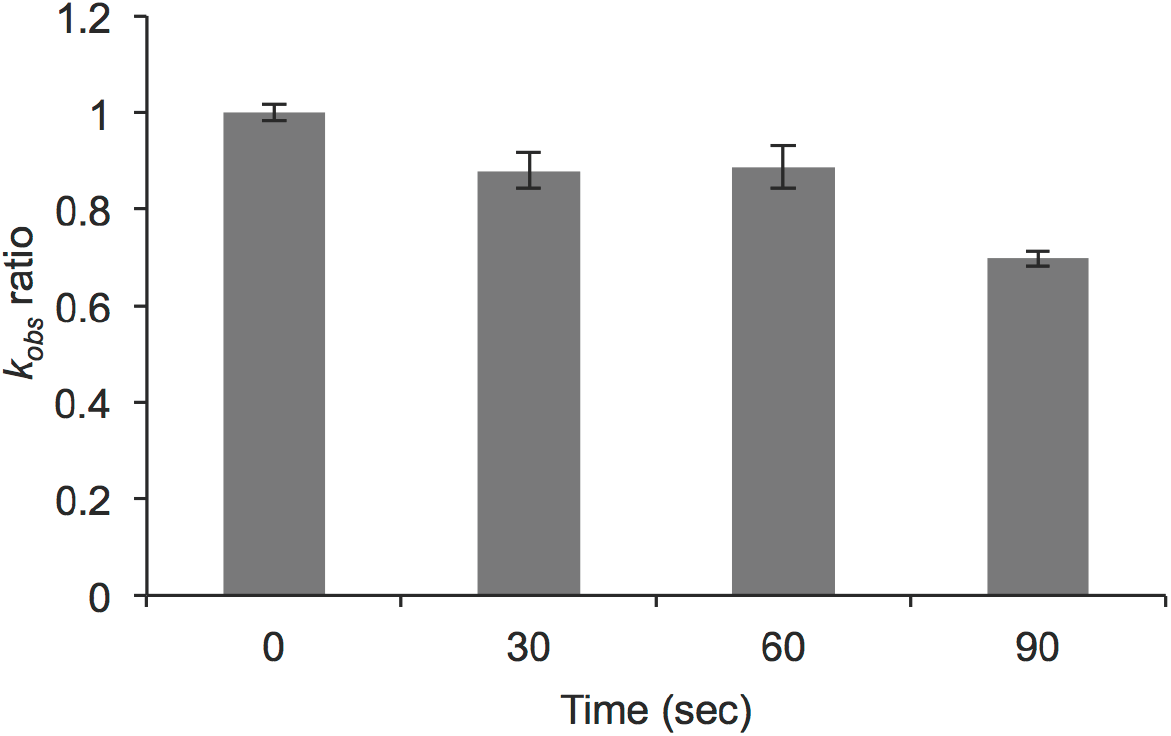
sCPR functionality at different time points during Copper(I)- catalyzed azide-alkyne cycloaddition (CuAAC) between sCPR_160PrF_ F and Sulfo- Cy3-Azide at 500 μM CuSO_4_. Functionality was determined by cytochrome ***c*** reduction efficiency and expressed as a fraction of the measured ***k*_*obs*_** at t=n over the ***k*_*obs*_** at t=0 and shows that only a minor loss of activity (around 10%) is observed up to 60 sec of reaction time at RT.

**Figure S7:**
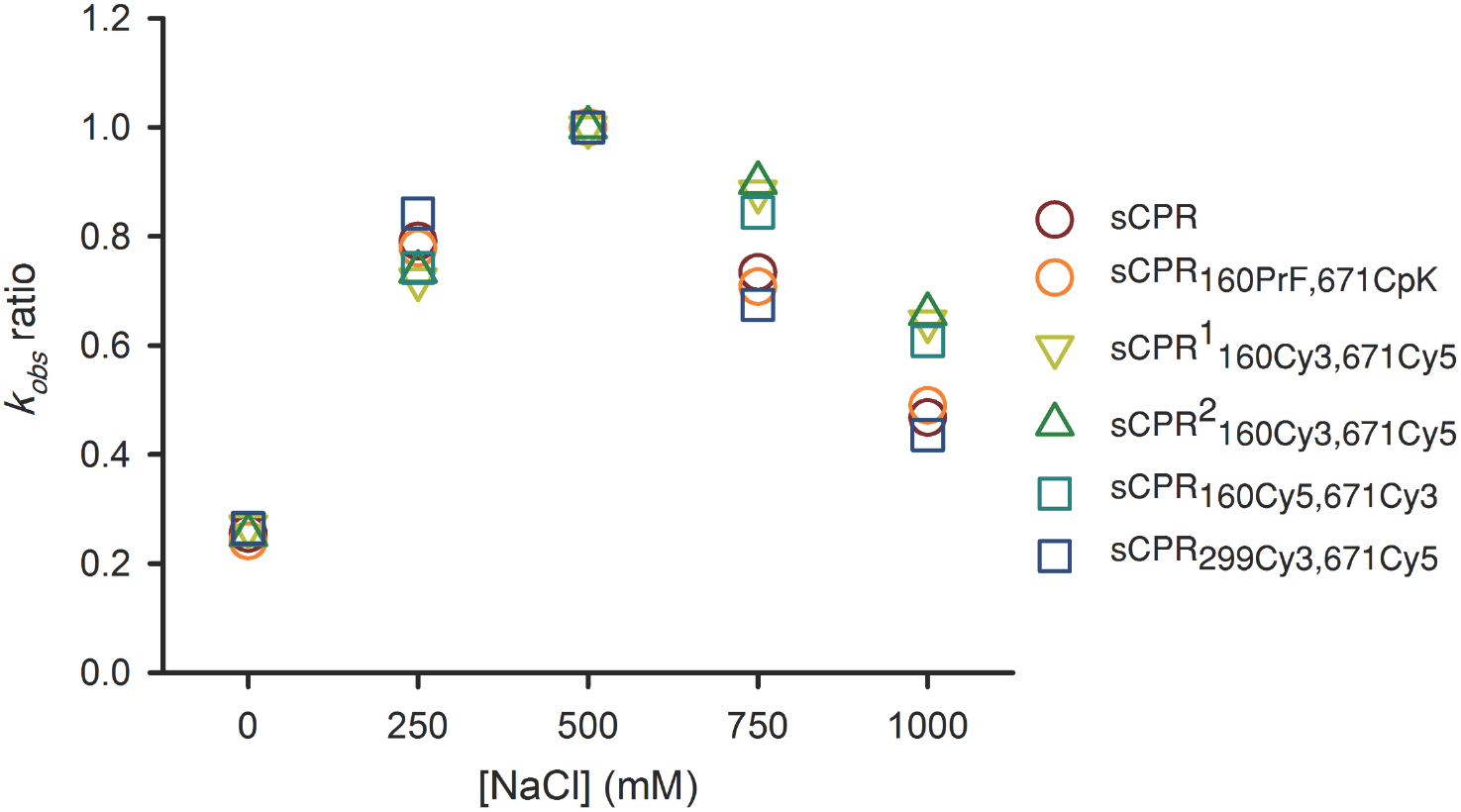
Salt-dependent cytochrome *c* reduction efficiency profile expressed as a ratio of the maximum *k*_*obs*_ value at 500 mM NaCl for each sCPR variant. All sCPR variants exhibit similarly shaped profiles in response to changing salt concentrations, which reflects the minor impact of labels on sCPR functionality. For explanations on the different mutants see main text.

**Figure S8:**
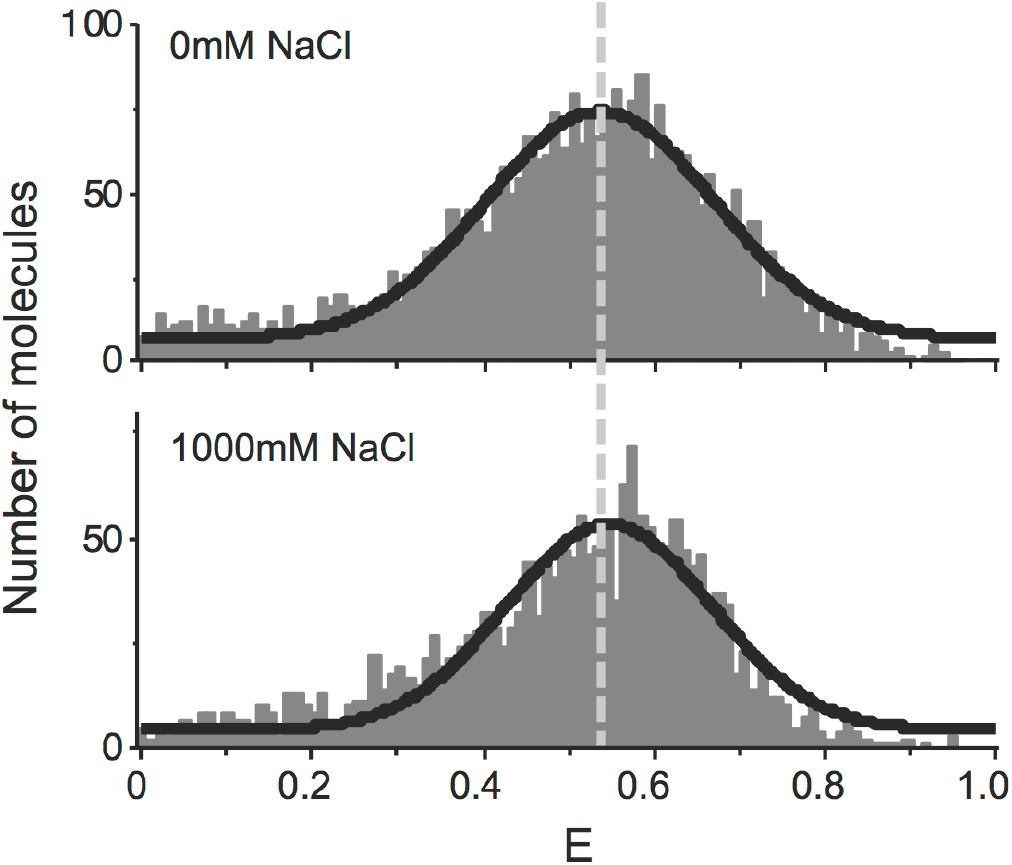
FRET efficiency histograms of sCPR_299Cy3,671Cy5_ at 0 and 1000 mM NaCl. No change in the shape and the peak value of the FRET efficiency are observed, thereby confirming that the presence of NaCl alone has no influence. As such, observed changes in the FRET efficiency are due to conformational changes, resulting in altered average distances between the two fluorophores.

### Supporting Tables

**Table S1:** **Quantum yields (QYs) used for correction of FRET efficiencies.** QYs were determined for singly labeled sCPR variants by the comparative method.

|  | Cy3 |  | Cy5 |  |
| --- | --- | --- | --- | --- |
|  | 0 NaCl | 1M NaCl | 0 NaCl | 1M NaCl |
| sCPR<sub>160Cy3,671Cy5</sub> | 0.15 | 0.18 | 0.16 | 0.15 |
| sCPR<sub>160Cy5,671Cy3</sub> | 0.18 | 0.17 | 0.17 | 0.2 |

**Table S2:** **Experimentally determined Forster radii R_0_ given in Angstrom.**

|  | 0 NaCl | 1M NaCl |
| --- | --- | --- |
| sCPR<sub>160Cy3,671Cy5</sub> | 55.3 | 57.5 |
| sCPR<sub>160Cy5,671Cy3</sub> | 56.1 | 56.5 |

**Table S3:** **Total average inter-dyes distances.** Distances were calculated from the corrected FRET efficiencies at different salt concentrations with sCPR_160Cy3,671Cy5_ in the presence and absence of 500 μM NADP+ (distances are displayed in Å).

| [NaCl] (M) | Average distance without NADP<sup>+</sup> | Average distance with NADP<sup>+</sup> |
| --- | --- | --- |
| 0.00 | 39.8 | 42.0 |
| 0.25 | 40.9 | 40.1 |
| 0.50 | 48.3 | 44.8 |
| 0.75 | 53.8 | 47.8 |
| 1.00 | 57.1 | 53.3 |

**Table S4:**
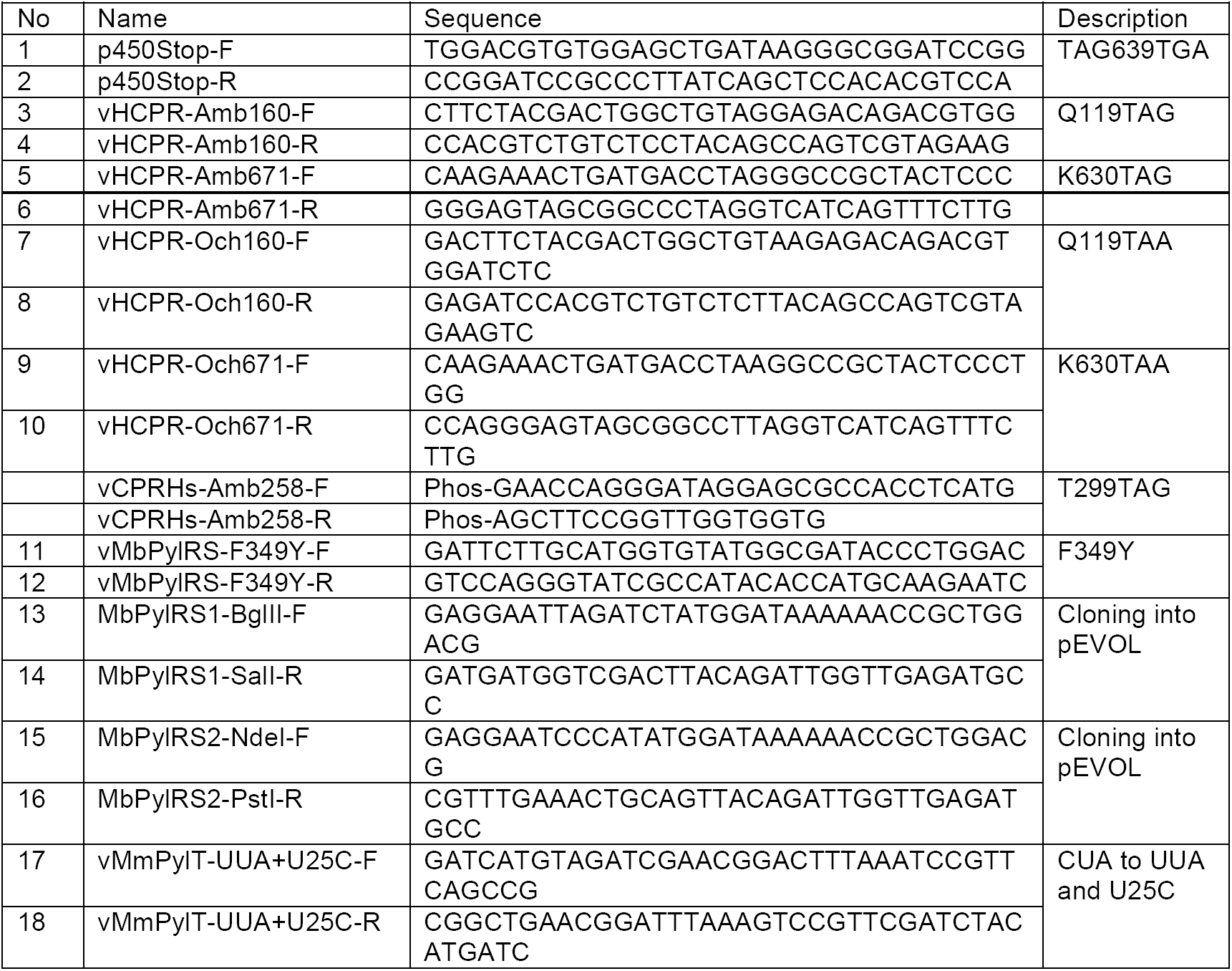
**Primer Sequences**

## Gene and protein sequences

Triplet codons and amino acids mutated in the course of this study are highlighted in red.

### sCPR

ATGGGCAGCAGCCATCATCATCATCATCACAGCAGCGGCCTGGTGCCGCGCGGCAGCCA

TATGCTCGAGTCCTCTGTCAGAGAGAGCAGCTTTGTGGAAAAGATGAAGAAAACGGGGAG

GAACATCATCGTGTTCTACGGCTCCCAGACGGGGACTGCAGAGGAGTTTGCCAACCGCCT

GTCCAAGGACGCCCACCGCTACGGGATGCGAGGCATGTCAGCGGACCCTGAGGAGTATG

ACCTGGCCGACCTGAGCAGCCTGCCAGAGATCGACAACGCCCTGGTGGTTTTCTGCATGG

CCACCTACGGTGAGGGAGACCCCACCGACAATGCCCAGGACTTCTACGACTGGCTGCAG

GAGACAGACGTGGATCTCTCTGGGGTCAAGTTCGCGGTGTTTGGTCTTGGGAACAAGACC

TACGAGCACTTCAATGCCATGGGCAAGTACGTGGACAAGCGGCTGGAGCAGCTCGGCGC

CCAGCGCATCTTTGAGCTGGGGTTGGGCGACGACGATGGGAACTTGGAGGAGGACTTCAT

CACCTGGCGAGAGCAGTTCTGGCCGGCCGTGTGTGAACACTTTGGGGTGGAAGCCACTG

GCGAGGAGTCCAGCATTCGCCAGTACGAGCTTGTGGTCCACACCGACATAGATGCGGCCA

AGGTGTACATGGGGGAGATGGGCCGGCTGAAGAGCTACGAGAACCAGAAGCCCCCCTTT

GATGCCAAGAATCCGTTCCTGGCTGCAGTCACCACCAACCGGAAGCTGAACCAGGGAACC

GAGCGCCACCTCATGCACCTGGAATTGGACATCTCGGACTCCAAAATCAGGTATGAATCTG

GGGACCACGTGGCTGTGTACCCAGCCAACGACTCTGCTCTCGTCAACCAGCTGGGCAAAA

TCCTGGGTGCCGACCTGGACGTCGTCATGTCCCTGAACAACCTGGATGAGGAGTCCAACA

AGAAGCACCCATTCCCGTGCCCTACGTCCTACCGCACGGCCCTCACCTACTACCTGGACA

TCACCAACCCGCCGCGTACCAACGTGCTGTACGAGCTGGCGCAGTACGCCTCGGAGCCC

TCGGAGCAGGAGCTGCTGCGCAAGATGGCCTCCTCCTCCGGCGAGGGCAAGGAGCTGTA

CCTGAGCTGGGTGGTGGAGGCCCGGAGGCACATCCTGGCCATCCTGCAGGACTGCCCGT

CCCTGCGGCCCCCCATCGACCACCTGTGTGAGCTGCTGCCGCGCCTGCAGGCCCGCTAC

TACTCCATCGCCTCATCCTCCAAGGTCCACCCCAACTCTGTGCACATCTGTGCGGTGGTTG

TGGAGTACGAGACCAAGGCTGGCCGCATCAACAAGGGCGTGGCCACCAACTGGCTGCGG

GCCAAGGAGCCTGCCGGGGAGAACGGCGGCCGTGCGCTGGTGCCCATGTTCGTGCGCA

AGTCCCAGTTCCGCCTGCCCTTCAAGGCCACCACGCCTGTCATCATGGTGGGCCCCGGCA

CCGGGGTGGCACCCTTCATAGGCTTCATCCAGGAGCGGGCCTGGCTGCGACAGCAGGGC

AAGGAGGTGGGGGAGACGCTGCTGTACTACGGCTGCCGCCGCTCGGATGAGGACTACCT

GTACCGGGAGGAGCTGGCGCAGTTCCACAGGGACGGTGCGCTCACCCAGCTCAACGTGG

CCTTCTCCCGGGAGCAGTCCCACAAGGTCTACGTCCAGCACCTGCTAAAGCAAGACCGAG

AGCACCTGTGGAAGTTGATCGAAGGCGGTGCCCACATCTACGTCTGTGGGGATGCACGGA

ACATGGCCAGGGATGTGCAGAACACCTTCTACGACATCGTGGCTGAGCTCGGGGCCATG

GAGCACGCGCAGGCGGTGGACTACATCAAGAAACTGATGACCAAGGGCCGCTACTCCCTG

GACGTGTGGAGCTGA

MGSSHHHHHHSSGLVPRGSHMLESSVRESSFVEKMKKTGRNIIVFYGSQTGTAEEFANRLSK

DAHRYGMRGMSADPEEYDLADLSSLPEIDNALVVFCMATYGEGDPTDNAQDFYDWLQETDVD

LSGVKFAVFGLGNKTYEHFNAMGKYVDKRLEQLGAQRIFELGLGDDDGNLEEDFITWREQFW

PAVCEHFGVEATGEESSIRQYELVVHTDIDAAKVYMGEMGRLKSYENQKPPFDAKNPFLAAVT

TNRKLNQGTERHLMHLELDISDSKIRYESGDHVAVYPANDSALVNQLGKILGADLDVVMSLNNL

DEESNKKHPFPCPTSYRTALTYYLDITNPPRTNVLYELAQYASEPSEQELLRKMASSSGEGKEL

YLSWVVEARRHILAILQDCPSLRPPIDHLCELLPRLQARYYSIASSSKVHPNSVHICAVVVEYETK

AGRINKGVATNWLRAKEPAGENGGRALVPMFVRKSQFRLPFKATTPVIMVGPGTGVAPFIGFI

QERAWLRQQGKEVGETLLYYGCRRSDEDYLYREELAQFHRDGALTQLNVAFSREQSHKVYV

QHLLKQDREHLWKLIEGGAHIYVCGDARNMARDVQNTFYDIVAELGAMEHAQAVDYIKKLMT

KGRYSLDVWS*

### *M.barkeri* PylRS

ATGGATAAAAAACCGCTGGACGTTCTGATCTCCGCTACGGGTCTGTGGATGAGCCGCACG

GGTACGCTGCATAAAATTAAACACCACGAAGTGTCACGTTCGAAAATCTATATCGAAATGG

CGTGCGGTGATCATCTGGTGGTTAACAATAGCCGTTCTTGTCGCACCGCGCGTGCCTTTC

GCCATCACAAATACCGCAAAACGTGCAAACGTTGTCGCGTGTCAGATGAAGACATTAACAA

TTTCCTGACCCGTAGTACGGAATCCAAAAACTCAGTGAAAGTTCGCGTCGTGAGTGCTCCG

AAAGTTAAAAAAGCGATGCCGAAAAGTGTCTCCCGTGCCCCGAAACCGCTGGAAAACTCA

GTGTCGGCAAAAGCTTCCACCAATACGAGCCGCTCTGTTCCGTCGCCGGCAAAAAGCACC

CCGAACAGCTCTGTCCCGGCAAGCGCACCGGCACCGTCTCTGACGCGTAGTCAGCTGGA

TCGCGTGGAAGCCCTGCTGTCCCCGGAAGACAAAATCTCACTGAATATGGCAAAACCGTTT

CGTGAACTGGAACCGGAACTGGTTACCCGTCGCAAAAACGATTTCCAACGTCTGTATACGA

ATGATCGCGAAGACTACCTGGGTAAACTGGAACGTGATATCACCAAATTTTTCGTGGACCG

CGGCTTTCTGGAAATCAAATCTCCGATTCTGATCCCGGCTGAATATGTTGAACGCATGGGT

ATTAACAATGATACCGAACTGAGTAAACAGATTTTTCGTGTGGATAAAAACCTGTGCCTGCG

GCCGATGCTGGCACCGACGCTGTATAATTACCTGCGTAAACTGGATCGCATTCTGCCGGG

TCCGATTAAAATCTTTGAAGTGGGCCCGTGTTATCGTAAAGAATCGGATGGCAAAGAACAC

CTGGAAGAATTTACCATGGTTAACTTCTGCCAAATGGGCAGCGGTTGTACGCGCGAAAATC

TGGAAGCGCTGATCAAAGAATTCCTGGATTACCTGGAAATCGACTTCGAAATCGTCGGTGAT

TCTTGCATGGTGTATGGCGATACCCTGGACATCATGCATGGTGACCTGGAACTGAGTTCCG

CTGTTGTCGGTCCGGTCAGCCTGGATCGTGAATGGGGCATTGACAAACCGTGGATCGGC

GCGGGTTTTGGCCTGGAACGCCTGCTGAAAGTTATGCACGGCTTCAAAAACATCAAACGT

GCGTCTCGCTCGGAATCGTATTACAACGGCATCTCAACCAATCTGTAA

MDKKPLDVLISATGLWMSRTGTLHKIKHHEVSRSKIYIEMACGDHLVVNNSRSCRTARAFRHHK

YRKTCKRCRVSDEDINNFLTRSTESKNSVKVRVVSAPKVKKAMPKSVSRAPKPLENSVSAKAS

TNTSRSVPSPAKSTPNSSVPASAPAPSLTRSQLDRVEALLSPEDKISLNMAKPFRELEPELVTR

RKNDFQRLYTNDREDYLGKLERDITKFFVDRGFLEIKSPILIPAEYVERMGINNDTELSKQIFRVD

KNLCLRPMLATLYNYLRKLDRILPGPIKIFEVGPCYRKESDGKEHLEEFTMVNFCQMGSGCTR

ENLEALIKEFLDYLEIDFEIVGDSCMVYGDTLDIMHGDLELSSAVVGPVSLDREWGIDKPWIGAG

FGLERLLKVMHGFKNIKRASRSESYYNGISTNL*

### Optimized *M. mazei* tRNA_UAA_

GGAAACCTGATCATGTAGATCGAACGGACTTTAAATCCGTTCAGCCGGGTTAGATTCCCGG GGTTTCCGCCA

### *M. janaschii* PrFRS

ATGGACGAGTTCGAAATGATTAAACGCAACACCAGCGAAATTATCTCTGAAGAAGAGCTGCG

CGAGGTGCTGAAGAAAGACGAGAAGAGCGCGGCGATTGGCTTTGAGCCGTCCGGTAAA

ATTCACCTGGGTCACTACCTGCAAATCAAGAAGATGATTGATCTGCAAAACGCTGGTTTTG

ACATCATTATCCTGCTGGCGGACCTGCACGCCTACCTGAATCAAAAGGGCGAGCTGGATG

AGATTCGCAAGATCGGCGACTACAATAAGAAAGTCTTCGAAGCCATGGGTTTGAAGGCTAA

ATACGTCTACGGTAGCCCGTTTCAGCTGGATAAGGATTACACGTTGAATGTGTACCGTCTG

GCGCTGAAAACCACGCTGAAACGCGCCCGTCGTTCCATGGAGCTGATTGCGCGCGAGGA

TGAGAATCCAAAAGTTGCTGAGGTTATTTACCCTATTATGCAAGTTAATGCTATTCACTACG

CGGGTGTTGATGTTGCCGTCGGTGGTATGGAGCAACGCAAAATTCACATGCTGGCACGTG

AACTGCTGCCGAAAAAGGTTGTCTGTATTCATAATCCGGTCCTGACCGGCCTGGATGGCG

AGGGTAAAATGAGCAGCAGCAAGGGTAACTTTATTGCAGTTGACGATAGCCCGGAAGAAAT

CCGTGCGAAGATCAAGAAAGCGTACTGCCCGGCAGGCGTGGTTGAGGGTAACCCGATCAT

GGAAATCGCCAAGTATTTTCTGGAATACCCACTGACGATTAAGCGCCCGGAGAAATTTGGC

GGCGACCTGACCGTCAACAGCTACGAGGAGCTGGAAAGCTTGTTTAAGAACAAAGAACTG

CATCCGATGCGCCTGAAAAACGCCGTGGCGGAAGAGCTGATTAAGATTCTGGAACCAATT

CGCAAACGTCTGTAA

MDEFEMIKRNTSEIISEEELREVLKKDEKSAAIGFEPSGKIHLGHYLQIKKMIDLQNAGFDIIILLAD

LHAYLNQKGELDEIRKIGDYNKKVFEAMGLKAKYVYGSPFQLDKDYTLNVYRLALKTTLKRARR

SMELIAREDENPKVAEVIYPIMQVNAIHYAGVDVAVGGMEQRKIHMLARELLPKKVVCIHNPVLTG

LDGEGKMSSSKGNFIAVDDSPEEIRAKIKKAYCPAGVVEGNPIMEIAKYFLEYPLTIKRPEKFG

GDLTVNSYEELESLFKNKELHPMRLKNAVAEELIKILEPIRKRL*

### *M. janaschii* tRNA_CUA_

CCGGCGGTAGTTCAGCAGGGCAGAACGGCGGACTCTAAATCCGCATGGCAGGGGTTCAAATCCCCTCCGCCGGA

## REFERENCES

1. Lerner, E.; Cordes, T.; Ingargiola, A.; Alhadid, Y.; Chung, S.; Michalet, X.; Weiss, S. Toward Dynamic Structural Biology: Two Decades of Single-Molecule Förster Resonance Energy Transfer. Science 2018, 359 (6373), eaan1133.

2. Pudney, C. R.; Khara, B.; Johannissen, L. O.; Scrutton, N. S. Coupled Motions Direct Electrons along Human Microsomal P450 Chains. PLoS Biol. 2011, 9 (12), e1001222.

3. Huang, W.-C.; Ellis, J.; Moody, P. C. E.; Raven, E. L.; Roberts, G. C. K. Redox-Linked Domain Movements in the Catalytic Cycle of Cytochrome P450 Reductase. Structure 2013, 21 (9), 1581–1589.

4. Laursen, T.; Singha, A.; Rantzau, N.; Tutkus, M.; Borch, J.; Hedegård, P.; Stamou, D.; Møller, B. L.; Hatzakis, N. S. Single Molecule Activity Measurements of Cytochrome P450 Oxidoreductase Reveal the Existence of Two Discrete Functional States. ACS Chem. Biol. 2014, 9 (3), 630–634.

5. Frances, O.; Fatemi, F.; Pompon, D.; Guittet, E.; Sizun, C.; Pérez, J.; Lescop, E.; Truan, G. A Well-Balanced Preexisting Equilibrium Governs Electron Flux Efficiency of a Multidomain Diflavin Reductase. Biophys. J. 2015, 108 (6), 1527–1536.

6. Kovrigina, E. A.; Pattengale, B.; Xia, C.; Galiakhmetov, A. R.; Huang, J.; Kim, J.-J. P.; Kovrigin, E. L. Conformational States of Cytochrome P450 Oxidoreductase Evaluated by Förster Resonance Energy Transfer Using Ultrafast Transient Absorption Spectroscopy. Biochemistry 2016, 55 (43), 5973–5976.

7. Riddick, D. S.; Ding, X.; Wolf, C. R.; Porter, T. D.; Pandey, A. V.; Zhang, Q.-Y.; Gu, J.; Finn, R. D.; Ronseaux, S.; McLaughlin, L. A.; et al. NADPH–Cytochrome P450 Oxidoreductase: Roles in Physiology, Pharmacology, and Toxicology. Drug Metab. Dispos. 2013, 41 (1), 12–23.

8. Hamdane, D.; Xia, C.; Im, S.-C.; Zhang, H.; Kim, J.-J. P.; Waskell, L. Structure and Function of an NADPH-Cytochrome P450 Oxidoreductase in an Open Conformation Capable of Reducing Cytochrome P450. J. Biol. Chem. 2009, 284 (17), 11374–11384.

9. Aigrain, L.; Pompon, D.; Moréra, S.; Truan, G. Structure of the Open Conformation of a Functional Chimeric NADPH Cytochrome P450 Reductase. EMBO Rep. 2009, 10 (7), 742–747.

10. Aramendia, P. F.; Negri, R. M.; Roman, E. S. Temperature Dependence of Fluorescence and Photoisomerization in Symmetric Carbocyanines.Influence of Medium Viscosity and Molecular Structure. J. Phys. Chem. 1994, 98 (12), 3165–3173.

11. Sadoine, M.; Cerminara, M.; Kempf, N.; Gerrits, M.; Fitter, J.; Katranidis, A. Selective Double-Labeling of Cell-Free Synthesized Proteins for More Accurate smFRET Studies. Anal. Chem. 2017, 89 (21), 11278–11285.

12. Chatterjee, A.; Sun, S. B.; Furman, J. L.; Xiao, H.; Schultz, P. G. A Versatile Platform for Single- and Multiple-Unnatural Amino Acid Mutagenesis in Escherichia coli. Biochemistry 2013, 52 (10), 1828–1837.

13. Kim, J.; Seo, M.-H.; Lee, S.; Cho, K.; Yang, A.; Woo, K.; Kim, H.-S.; Park, H.-S. Simple and Efficient Strategy for Site-Specific Dual Labeling of Proteins for Single-Molecule Fluorescence Resonance Energy Transfer Analysis. Anal. Chem. 2013, 85 (3), 1468–1474.

14. Sachdeva, A.; Wang, K.; Elliott, T.; Chin, J. W. Concerted, Rapid, Quantitative, and Site-Specific Dual Labeling of Proteins. J. Am. Chem. Soc. 2014, 136 (22), 7785–7788.

15. Wang, K.; Sachdeva, A.; Cox, D. J.; Wilf, N. M.; Lang, K.; Wallace, S.; Mehl, R. A.; Chin, J. W. Optimized Orthogonal Translation of Unnatural Amino Acids Enables Spontaneous Protein Double-Labelling and FRET. Nat. Chem. 2014, 6 (5), 393–403.

16. Shen, A. L.; Christensen, M. J.; Kasper, C. B. NADPH-Cytochrome P-450 Oxidoreductase. The Role of Cysteine 566 in Catalysis and Cofactor Binding. J. Biol. Chem. 1991, 266 (30), 19976–19980.

17. Deiters, A.; Schultz, P. G. In Vivo Incorporation of an Alkyne into Proteins in Escherichia coli. Bioorg. Med. Chem. Lett. 2005, 15 (5), 1521–1524.

18. Dietrich, A.; Buschmann, V.; Müller, C.; Sauer, M. Fluorescence Resonance Energy Transfer (FRET) and Competing Processes in Donor-Acceptor Substituted DNA Strands: A Comparative Study of Ensemble and Single-Molecule Data. J. Biotechnol. 2002, 82 (3), 211–231.

19. Young, T. S.; Ahmad, I.; Yin, J. A.; Schultz, P. G. An Enhanced System for Unnatural Amino Acid Mutagenesis in E. coli. J. Mol. Biol. 2010, 395 (2), 361–374.

20. Campelo, D.; Lautier, T.; Urban, P.; Esteves, F.; Bozonnet, S.; Truan, G.; Kranendonk, M. The Hinge Segment of Human NADPH-Cytochrome P450 Reductase in Conformational Switching: The Critical Role of Ionic Strength. Front. Pharmacol. 2017, 8.

21. Olofsson, L.; Margeat, E. Pulsed Interleaved Excitation Fluorescence Spectroscopy with a Supercontinuum Source. Opt. Express 2013, 21 (3), 3370–3378.

22. Sanborn, M. E.; Connolly, B. K.; Gurunathan, K.; Levitus, M. Fluorescence Properties and Photophysics of the Sulfoindocyanine Cy3 Linked Covalently to DNA. J. Phys. Chem. B 2007, 111 (37), 11064–11074.

23. Berlier, J. E.; Rothe, A.; Buller, G.; Bradford, J.; Gray, D. R.; Filanoski, B. J.; Telford, W. G.; Yue, S.; Liu, J.; Cheung, C.-Y.; et al. Quantitative Comparison of Long-Wavelength Alexa Fluor Dyes to Cy Dyes: Fluorescence of the Dyes and Their Bioconjugates. J. Histochem. Cytochem. Off. J. Histochem. Soc. 2003, 51 (12), 1699–1712.

24. Bavishi, K.; Li, D.; Eiersholt, S.; Hooley, E. N.; Petersen, T. C.; Møller, B. L.; Hatzakis, N. S.; Laursen, T. Direct Observation of Multiple Conformational States in Cytochrome P450 Oxidoreductase and Their Modulation by Membrane Environment and Ionic Strength. Sci. Rep. 2018, 8 (1).

25. Ellis, J.; Gutierrez, A.; Barsukov, I. L.; Huang, W.-C.; Grossmann, J. G.; Roberts, G. C. K. Domain Motion in Cytochrome P450 Reductase: Conformational Equilibria Revealed by NMR and Small-Angle X-Ray Scattering. J. Biol. Chem. 2009, 284 (52), 36628–36637.

26. Wadsater, M.; Laursen, T.; Singha, A.; Hatzakis, N. S.; Stamou, D.; Barker, R.; Mortensen, K.; Feidenhans’l, R.; Lindberg Moller, B.; Cardenas, M. Monitoring Shifts in the Conformation Equilibrium of the Membrane Protein Cytochrome P450 Reductase (POR) in Nanodiscs. J. Biol. Chem. 2012.

27. Lu, W.; Wang, L.; Chen, L.; Hui, S.; Rabinowitz, J. D. Extraction and Quantitation of Nicotinamide Adenine Dinucleotide Redox Cofactors. Antioxid. Redox Signal.2018, 28 (3), 167–179.

28. Hedison, T. M.; Hay, S.; Scrutton, N. S. Real-Time Analysis of Conformational Control in Electron Transfer Reactions of Human Cytochrome P450 Reductase with Cytochrome c. FEBS J. 2015, 282 (22), 4357–4375.

29. Hay, S.; Brenner, S.; Khara, B.; Quinn, A. M.; Rigby, S. E. J.; Scrutton, N. S. Nature of the Energy Landscape for Gated Electron Transfer in a Dynamic Redox Protein. J. Am. Chem. Soc. 2010, 132 (28), 9738–9745.

## References

(1) Aigrain, L.; Pompon, D.; Moréra, S.; Truan, G. Structure of the Open Conformation of a Functional Chimeric NADPH Cytochrome P450 Reductase. EMBO Rep. 2009, 10 (7), 742–747.

(2) Young, T. S.; Ahmad, I.; Yin, J. A.; Schultz, P. G. An Enhanced System for Unnatural Amino Acid Mutagenesis in E. Coli. J. Mol. Biol. 2010, 395 (2), 361–374.

(3) Chatterjee, A.; Sun, S. B.; Furman, J. L.; Xiao, H.; Schultz, P. G. A Versatile Platform for Single- and Multiple-Unnatural Amino Acid Mutagenesis in Escherichia Coli. Biochemistry (Mosc.) 2013, 52 (10), 1828–1837.

(4) Deiters, A.; Schultz, P. G. In Vivo Incorporation of an Alkyne into Proteins in Escherichia Coli. Bioorg. Med. Chem. Lett. 2005, 15 (5), 1521–1524.

(5) Würth, C.; Grabolle, M.; Pauli, J.; Spieles, M.; Resch-Genger, U. Relative and Absolute Determination of Fluorescence Quantum Yields of Transparent Samples. Nat. Protoc. 2013, 8, 1535.

(6) Olofsson, L.; Felekyan, S.; Doumazane, E.; Scholler, P.; Fabre, L.; Zwier, J. M.; Rondard, P.; Seidel, C. A. M.; Pin, J.-P.; Margeat, E. Fine Tuning of Sub-Millisecond Conformational Dynamics Controls Metabotropic Glutamate Receptors Agonist Efficacy. Nat. Commun. 2014, 5, 5206.

(7) Widengren, J.; Kudryavtsev, V.; Antonik, M.; Berger, S.; Gerken, M.; Seidel, C. A. M. Single-Molecule Detection and Identification of Multiple Species by Multiparameter Fluorescence Detection. Anal. Chem. 2006, 78 (6), 2039–2050.

(8) Lee, N. K.; Kapanidis, A. N.; Wang, Y.; Michalet, X.; Mukhopadhyay, J.; Ebright, R. H.; Weiss, S. Accurate FRET Measurements within Single Diffusing Biomolecules Using Alternating-Laser Excitation. Biophys. J. 2005, 88 (4), 2939–2953.

(9) Hellenkamp, B.; Schmid, S.; Doroshenko, O.; Opanasyuk, O.; Kühnemuth, R.; Rezaei Adariani, S.; Ambrose, B.; Aznauryan, M.; Barth, A.; Birkedal, V.; et al. Precision and Accuracy of Single-Molecule FRET Measurements—a Multi-Laboratory Benchmark Study. Nat. Methods 2018, 15 (9), 669–676.

(10) McCammon, K. M.; Panda, S. P.; Xia, C.; Kim, J.-J. P.; Moutinho, D.; Kranendonk, M.; Auchus, R. J.; Lafer, E. M.; Ghosh, D.; Martasek, P.; et al. Instability of the Human Cytochrome P450 Reductase A287P Variant Is the Major Contributor to Its Antley-Bixler Syndrome-like Phenotype. J. Biol. Chem. 2016, 291 (39), 20487–20502.

(11) Frances, O.; Fatemi, F.; Pompon, D.; Guittet, E.; Sizun, C.; Pérez, J.; Lescop, E.; Truan, G. A Well-Balanced Preexisting Equilibrium Governs Electron Flux Efficiency of a Multidomain Diflavin Reductase. Biophys. J. 2015, 108 (6), 1527–1536.

